# *Wilms Tumor 1b* defines a wound-specific sheath cell subpopulation associated with notochord repair

**DOI:** 10.1101/172288

**Authors:** Juan Carlos Lopez-Baez, Zhiqiang Zeng, Witold Rybski, Leonie F.A. Huitema, Alessandro Brombin, Rodney M. Dale, Koichi Kawakami, Christoph Englert, Stefan Schulte-Merker, Nicholas D. Hastie, E. Elizabeth Patton

## Abstract

Regenerative therapy for degenerative spine disorders requires the identification of cells that can slow down and possibly reverse degenerative processes. Here, we identify a novel and unanticipated wound-specific notochord sheath cell subpopulation that expresses Wilms Tumor (WT) 1b following injury. Using live imaging in zebrafish, we show that localized damage leads to Wt1b expression in the sheath, and that *wt1b*+ cells migrate into the wound to form a stopper-like structure, likely to maintain structural integrity. At the wound *wt1b+* and *entpd5*+ cells constitute distinct subpopulations, and mark the site of an extra vertebra that forms in an untypical manner via a cartilage intermediate. Surprisingly, *wt1b*+ cells become closely associated with the chordacentra and sustain *wt1b* expression for over 35 days during vertebra formation. Given that remnants of notochord cells remain in the adult intervertebral disc, the identification of novel subpopulations may have important implications for regenerative treatments for spine disorders.

**Highlights:**

- Notochord injury triggers wound-specific expression of *wt1b* in novel sheath subpopulation
- WT1b notochord sheath cells fill injury site and form stopper-like structure
- WT1b subpopulation marks site of a new vertebra that forms via a cartilage intermediate
- WT1b wound-specific subpopulation perdures throughout and after vertebra repair

## Introduction

Wilms’ tumour 1 (WT1) is a zinc finger transcription factor that regulates key developmental stages of several mesodermal tissues including the kidneys, gonads and coronary vasculature (Hastie 2017). In the developing kidney, WT1 is required for the maintenance of mesenchymal nephron progenitors (Kriedberg et al, 1993, Motamedi et al, 2014) as well as differentiation of these progenitors into the epithelial components of the nephron (Essafi et al, 2011). In contrast, in the developing heart, WT1 is expressed in the epicardium (mesothelial lining) and required for the production, via an epithelial to mesenchymal transition (EMT), of coronary vascular progenitors (EPDCs) that migrate into the myocardium (Martinez-Estrada et al., 2010). Similarly, WT1-expressing mesothelium is the source of mesenchymal progenitors for specialised cell types within several other developing organs. These include stellate cells within the liver (Asahina et al, 2008), interstitial cells of Cajal in the intestine (Carmona et al, 2013) and adipocytes within visceral fat depots (Chau et al, 2014). WT1 expression is down-regulated in the epicardium postnatally but reactivated in response to tissue damage in both mice (Smart et al., 2011) and zebrafish (Schnabel et al., 2011). In both organisms, this activation of WT1 in response to damage is associated with new rounds of epicardial EMT, leading to the production of coronary vascular progenitors (Smart et al., 2011; Schnabel et al., 2011).

Given the reactivation of *Wt1/wt1b* in the damaged epicardium we set out to investigate whether WT1 programmes are initiated in response to other sources of tissue damage in zebrafish, and uncovered a novel Wt1 response to wounding of the notochord. The notochord is a transient embryonic structure that provides axial support, signalling information, and is required for vertebrae development and formation (Stemple et al., 2005). The notochord is comprised of two cell populations, the inner vacuolated cells that provide rigid support to the embryo, and the outer sheath cells, a single cell epithelial layer that surrounds the vacuolated cells and secretes components of the extracellular matrix to provide turgor pressure to the vacuolated cells (Ellis *et al*., 2013; Apschner et al., 2011). This rigid axial structure eventually is replaced by vertebrae bone. In zebrafish, a row of metameric mineralized rings, known as chordacentra, forms around the notochord in an anterior to posterior fashion and constitute the first signs of the definitive vertebrae. The chordacentra delineate the future sites where mature vertebra will form and ossify as the larva grows, while the notochord cells develop into the nucleus pulposus of the adult intervertebral disk, a soft gel-like tissue that provides cushioning and flexibility for the spine.

Degeneration of the intervertebral disk leads to extensive back pain, one of the top global causes of years lived with disability (Lawson & Harfe, 2015). Treatment primarily consists of managing the pain symptoms, or in more progressed disease includes extensive surgery. One of the major goals of the tissue-engineering field is to identify cells and tissues that will enable novel regenerative therapies to slow down and possibly reverse the degenerative process. Here, we uncover a novel cellular subpopulation in the notochord sheath that emerges at the site of damage and is maintained until formation of a repaired adult vertebra structure. Surprisingly, this subpopulation expresses *wt1b* despite no evidence of *wt1b* expression in physiological notochord development or ossification. Our findings suggest that the zebrafish notochord is protected by a novel wound-specific programme that seals the notochord wound in the embryo and establishes the site of a new adult vertebra.

## Results

### Wound specific expression of *wt1b* in the notochord

Given the expression of *wt1b* in the regenerating heart, we wanted to explore the expression of *wt1* in other regenerating tissues, and began with the tail fin regenerative processes. There are two *wt1* paralogues in zebrafish, *wt1a* and *wt1b*, and so we performed tail fin amputations on zebrafish larvae 3 days post fertilization (dpf) using *Tg(wt1a:GFP)* and *Tg(wt1b:GFP)* transgenic lines (Bollig *et al*., 2006; Perner *et al*., 2007; **Supplementary Figure 1a**). Surprisingly, we discovered that tail fin amputations that included partial removal of the notochord triggered a change of cellularity in the notochord, coupled with the specific, de *novo* upregulation of GFP in a *Tg(wt1b:GFP)* transgenic line. This response was specific to *wt1b* because we did not observe expression of GFP in *Tg(wt1a:GFP)* tail fin amputated larvae (**Supplementary 1b-e)**.

Next, we developed a needle-based assay to specifically induce localized damage in the developing zebrafish notochord independent of tail fin amputation. Needle injury was induced in 3 dpf *Tg(wt1b:GFP)* that had been crossed with *casper* fish to remove pigmentation and imaged at 72 hours post injury (hpi) (**Figure 1a**). Needle induced wounds triggered a similar, albeit stronger *wt1b:GFP* response to the tail fin amputations, that was specifically localised to the site of the wound (**Figure 1b**). Time course imaging showed a progressive expansion of the damaged area over 72 hours, with an increasing expression of GFP signal, concomitant with a change of cellularity in the notochord (**Figure 1c**). Importantly, this was not observed in uninjured zebrafish controls (**Figure 1c**) or in notochord injured *Tg(wt1a:GFP)* transgenic larvae (data not shown). Histological staining of the damaged area revealed the presence of a subpopulation of cells at the site of injury, which contrasted morphologically with the uniform, vacuolated inner cells of the notochord (**Figure 1d**). These cells stained positively for GFP and for endogenous Wt1 protein by immunohistochemistry, validating the faithful expression of the transgene with endogenous *wt1b* expression in this response (**Figure 1e**). Thus, following notochord injury, an unanticipated expression of *wt1b* marks a subpopulation of cells that emerges in the notochord and is associated with the wound.

**Figure 1.**
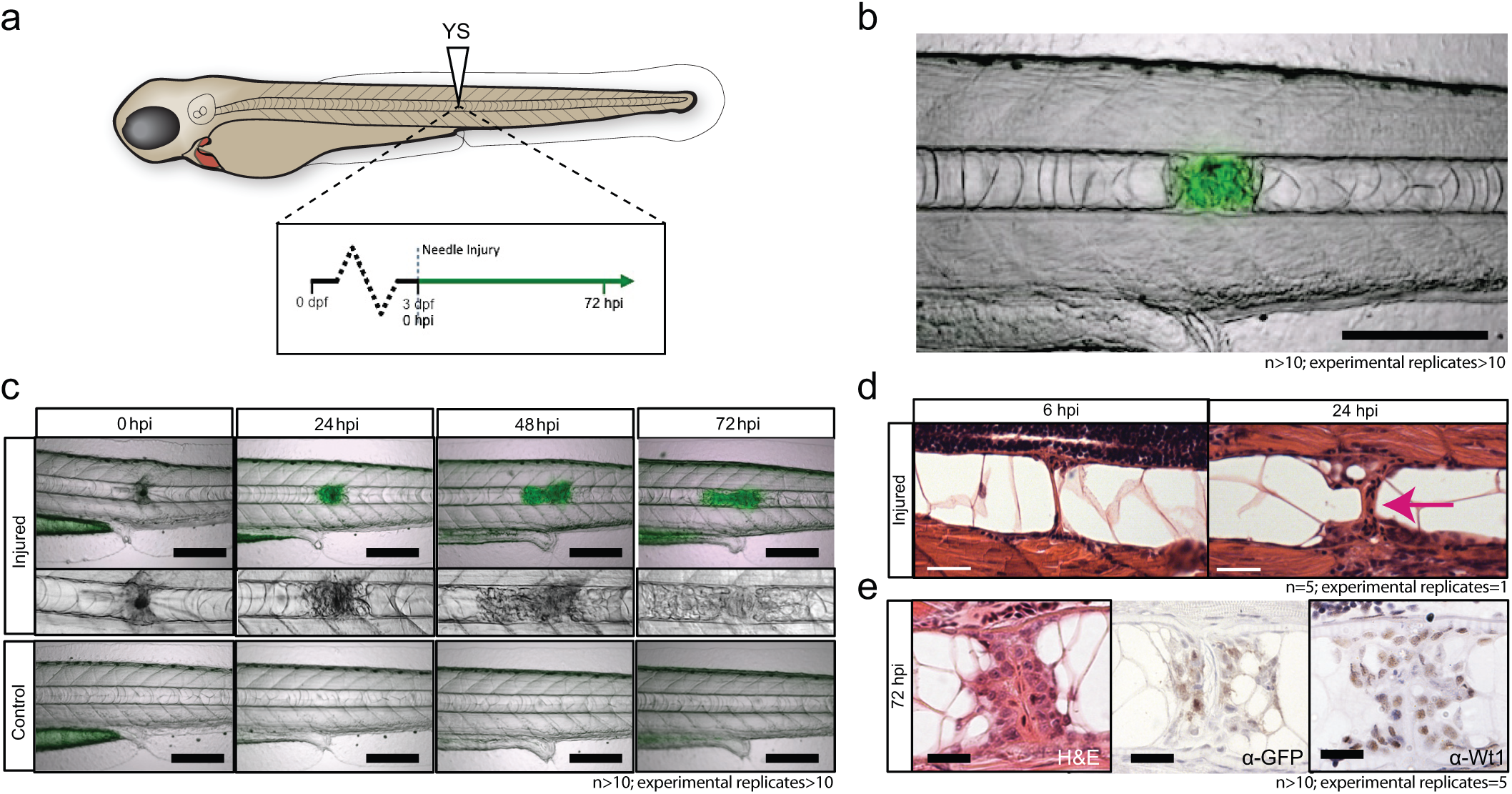
Notochord injury triggers a local and sustained expression Wt1. (a) Schematic of notochord needle-injury protocol. 3 dpf Tg(*wt1b:GFP;casper*) larvae are injured above the yolk sac (YS) and followed for 72 hours. (b, c) Images of Tg(*wt1b:GFP; casper*) zebrafish trunk over time following notochord needle injury, and uninjured matched controls. GFP signal is associated with a change of cellularity in the injured notochord (inset). n>10; experimental replicates>10. Scale bar: 100μm. (d) H&E staining of the injured area at 6 hpi and 24 hpi highlighted the progressive change in cellularity at the site of the injury (arrow). n=5; experimental replicates=1. Scale bar: 20μm. (e) Immunohistochemistry of the injured area with α-GFP and α-Wt1 antibodies. n>10; experimental replicates=5. Scale bar: 2μm. dpf = days post fertilization; hpi = hours post injury; H&E= haematoxylin and eosin.

### *wt1b* expressing cells emerge from the notochord sheath

To determine the origin of the wound-specific *wt1b*+ cells, we examined *wt1b* expression in the notochord and vacuolated cells, using a *Tg(SAGFF218:GFP)* transgenic line that labels the membrane of the inner vacuolated cells and *Tg(col2a1a:RFP)* that is specifically expressed in the surrounding notochord sheath cells (**Figure 2a;** Yamamoto *et al*., 2010 and Dale and Topczewski, 2011).

**Figure 2.**
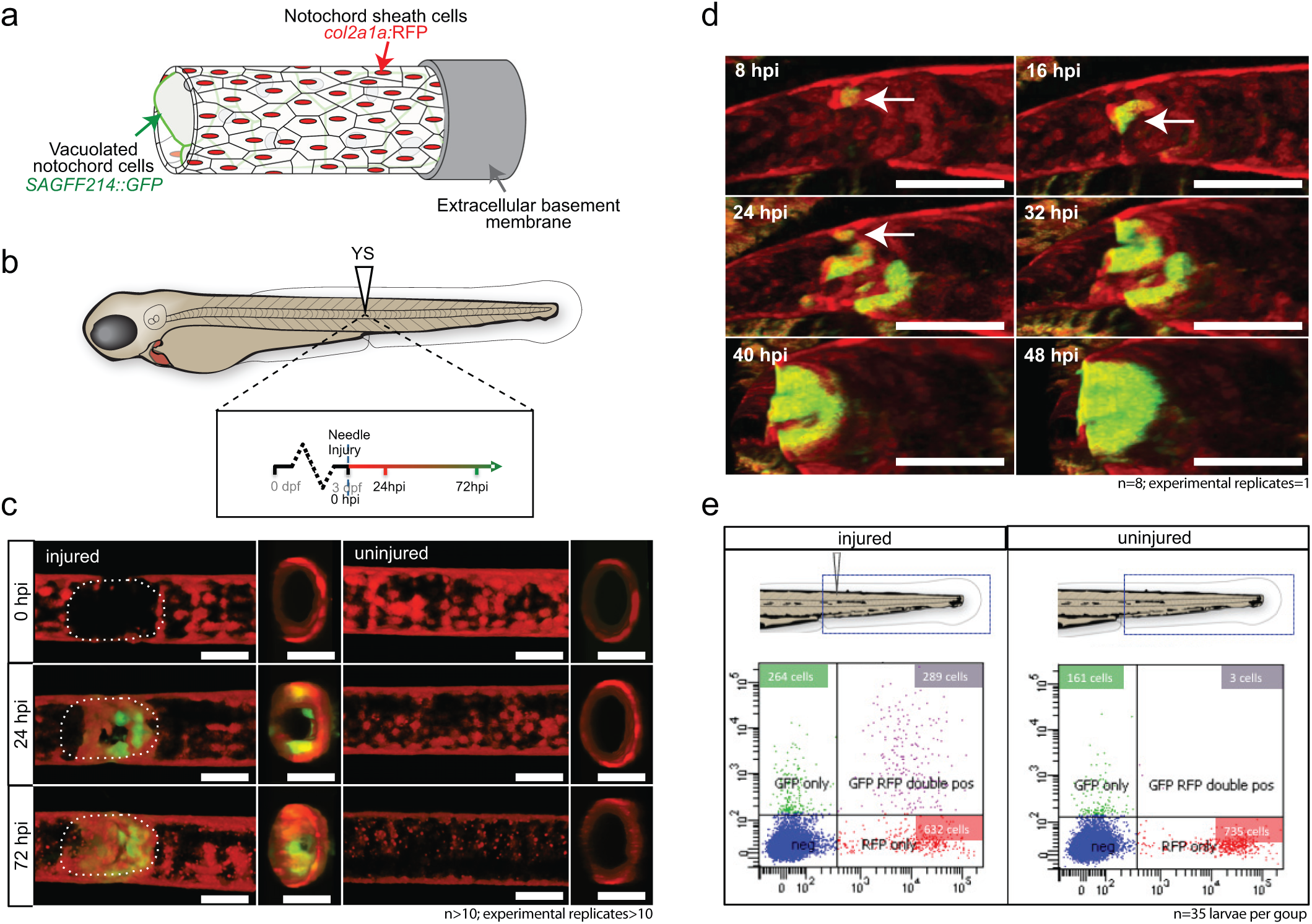
*wt1b:GFP* expressing notochord sheath cells populate the site of injury in the damaged notochords. (a) Schematic diagram of the notochord and transgenic lines used in this study. The notochord is composed of an inner population of highly vacuolated cells (green arrow), surrounded by a layer of epithelial-like sheath cells (red arrow), encapsulated by a thick layer of extracellular basement membrane (grey arrow). (b) Schematic of experimental design: 3dpf Tg(*wt1b:GFP;col2a1a:RFP;casper*) larvae were needle-injured and imaged at 0, 24 and 72 hpi. (c) Needle damage led to the formation of a cell-less gap in the layer of notochord sheath cells (0 hpi – injured; dashed line). GFP expression can be observed in the notochord sheath cells surrounding the area of damage by 24 hpi (inset) and these appear to engulf the injured area by 72 hpi (inset). n >10; experimental replicates > 10. Scale bar: 100μm. (d) Multiphoton time-lapse imaging of wound site. Initial upregulation of GFP occurs at 8 hpi in the *col2a1a:RFP* positive cells (arrow) and propagates across the injured area over the next 40 hours. n = 8; experimental replicates = 1. Scale bar: 100μm. (e) FACS analysis of cell populations in injured and non-injured zebrafish trunk tissue. GFP+RFP+ double positive cells are present in injured Tg(*wt1b:GFP;col2a1a:RFP*) at 72 hpi (n=35 larvae per group). dpf = days post fertilization; hpi = hours post injury.

A needle-induced notochord wound in the *Tg(SAGFF214:GFP)* transgenic line showed that GFP-expressing cells were lost rapidly upon injury, creating a gap in the row of vacuolated cells. Eventually, this gap was filled with new cells by 144 hpi (**Supplementary Figure 2a,b**). The *SAGFF214:GFP* response was distinct from the *wt1b*+ response in time (emerging at 72 hpi compared with 24 hpi), size and number (few and large compared with numerous and small), and in coverage of the wound (visible gaps remaining at the site compared with filling the damage site). These data suggest that *wt1b* expressing cells are distinct from the vacuolated cells at the site of injury.

Next, we explored the role of the notochord sheath cells in this process. We crossed the *Tg(wt1b:GFP)* transgenic line to a *Tg(col2a1a:RFP)* transgenic line, generated with a 1 KB fragment of the *col2a1a* promoter that is transiently expressed in the sheath cells until approximately 6 dpf (Dale & Topczewski, 2011). Live confocal and multiphoton imaging revealed *wt1b:GFP* expression in the *col2a1a:RFP* notochord sheath cells following needle induced notochord damage (**Figure 2 b-d**; **Video 1; Supplementary Figure 2c,d**). *Wt1b:GFP* co-expression with *col2a1a:RFP* was visible by 24 hpi in a ring surrounding the notochord vacuolated cells, and by 72 hpi the *wt1b:GFP* subpopulation of sheath cells had migrated into the inner lumen of the notochord to fill the wound and produce a visible stopper-like seal in the notochord.

To validate the co-expression of *wt1b:GFP* and *col2a1a:RFP* in the wounded fish, we FACS sorted cell populations in the injured versus uninjured larvae isolated from the trunk region (**Figure 2e**; n = 35 larvae per set). While GFP+ only and RFP+ only expressing cells were found in both injured and non-injured larvae, only the wounded fish had cells that co-expressed *wt1b:GFP* and *col2a1a:RFP* (GFP+RFP+; 289 cells vs. 3 cells respectively).

Our evidence indicates that the notochord wound triggers a unique *wt1b*+ subpopulation to emerge in the notochord sheath cells. This *wt1b*+ sheath cell subpopulation migrates into the wound and generates a stopper-like structure, possibly to prevent further loss of notochord turgor pressure and maintain notochord integrity.

### Notochord wounds express cartilage and mesenchyme genes

To address the molecular process at the site of the wound, we compared the transcriptome of the trunk region in the injured and uninjured 72 hpi larvae (**Figure 3a, b**; n = 50 larvae per subset). Microarray analysis revealed a highly significant 131-fold increase in expression of *matrix gla protein* (mgp), a gene that is known to express in chondrocytic zebrafish tissues (Gavaia *et al*., 2006) and to be involved in the inhibition of hydroxyapatite production during ectopic bone formation (Zebboudj *et al*., 2002; Sweatt *et al*., 2003; Schurgers *et al*., 2013) (**Figure 3c, d**). Other genes included mesenchymal and cell adhesion markers, such as *fn1b*, coagulation factors, such as *f13a1b*, and immune response genes, such as zgc:92041 and complement c6 (**Figure 3d**).

**Figure 3.**
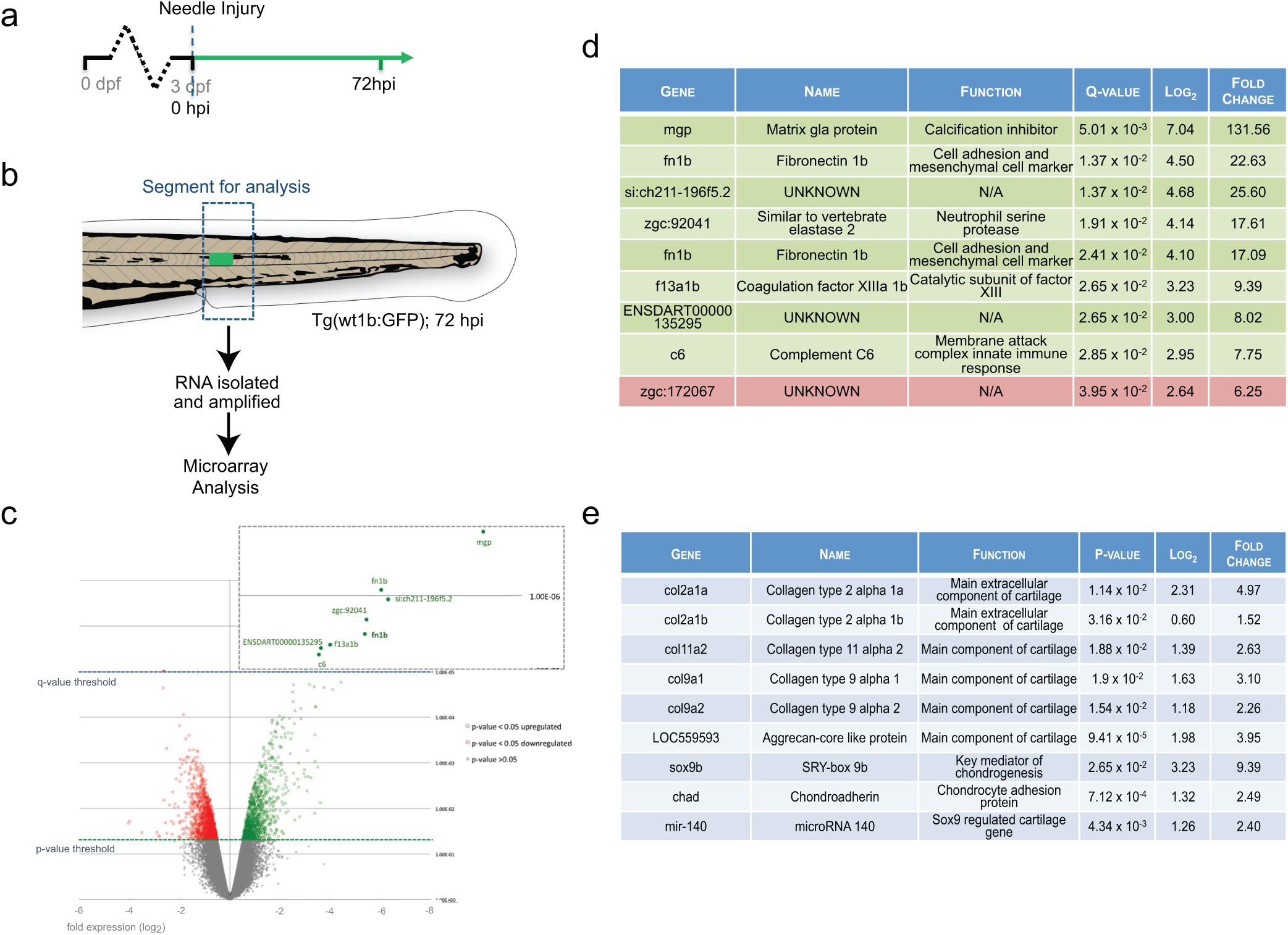
Cartilage genes are expressed in the notochord-injured zebrafish. (a) Experimental plan: 3 dpf Tg*(wt1b:GFP)* larvae were needle injured and grown for 72 hours with uninjured age-matched controls (n = 50 larvae per group). (b) Schematic of the area around the wt1b:GFP expression was excised at 72 hpi (dotted area) and RNA was extracted and amplified. A similar area was taken from age-matched uninjured controls. (c) Volcano plot displaying the differentially expressed genes between injured and non-injured larvae. The y-axis measures the mean expression value of log 10 (p-value) and separates upregulated from downregulated genes. The x-axis represents the log2 fold change of expression. Significantly upregulated genes are shown as green circles or dots and downregulated genes are shown as red circles or dots. Green dotted line represents the p-value threshold (p < 0.05) and blue dotted line represents the false discovery rate (FDR) or q-value threshold (q < 0.05). Genes with highest expression change in magnified view. (d) Table showing the most significantly differentially expressed genes in injured larvae (q < 0.05). Upregulated genes are shown in green and downregulated genes are shown in red. (e) Table showing cartilage-associated genes that were significantly upregulated in the injured larvae (p < 0.05).

The increased expression of *mgp* and *f13a1b* genes implicated the *de novo* acquisition of chondrogenic features in the injured tissues. Chondrogenic cells in the endochondral tissues of the craniofacial, fin bud and axial skeletons express *mgp* (Gavaia *et al*., 2006) and *FXIIIA* expression is localized to the developing chondrogenic mesenchyme of the pectoral fin bud (Deasey et al., 2012). The expression of cartilage genes was unexpected because ossification of the zebrafish notochord occurs via the formation the chordacentra, and does not require the establishment of cartilage anlagen (Flemming 2004; Bensimon-Briti et al., 2012; Lefebve & Bhattaram, 2010). To examine the expression of other chondrogenic genes, we analyzed the top 100 significant genes and found an increase in expression in Sox9, the master regulator of chondrogenesis, five collagen genes associated with chondrogenic tissues(*col2a1a*, *col2a1b*, *col11a2*, *col9a1* and *col9a2*), the cartilage specific extracellular structural protein Aggrecan, a microRNA regulator of chondrogenesis microRNA140 and the matrix-cell anchor protein chondroadherin (*chad*) (**Figure 3e**). Our results reveal that notochord wounding leads to an unexpected increase in expression of genes associated with cartilage.

### Extra vertebra forms at the repair site via an unusual cartilage intermediate

The expression of cartilage genes suggests that the notochord wound may induce a previously unknown and alternative bone development process. We stained injured and control animals with alcian blue and alizarin red stains, which highlight the cartilage and bone respectively. Cartilage was clearly visible at the site of injury as soon as 3 dpi. This staining was significantly stronger and distinct from the highly coordinated segmental cartilage staining that normally occurs during zebrafish vertebra development, which is clearly visible in both injured and non-injured controls by 14 dpi (**Figure 4a**). Similarly, the alizarin red dye identified the anterior to posterior forming chordacentra rings during larval development. However, in injured zebrafish larvae, the normally uniform mineralization pattern was interrupted around the site of damage, leading to delayed formation of the chordacentra at later stages (**Figure 4a**).

**Figure 4.**
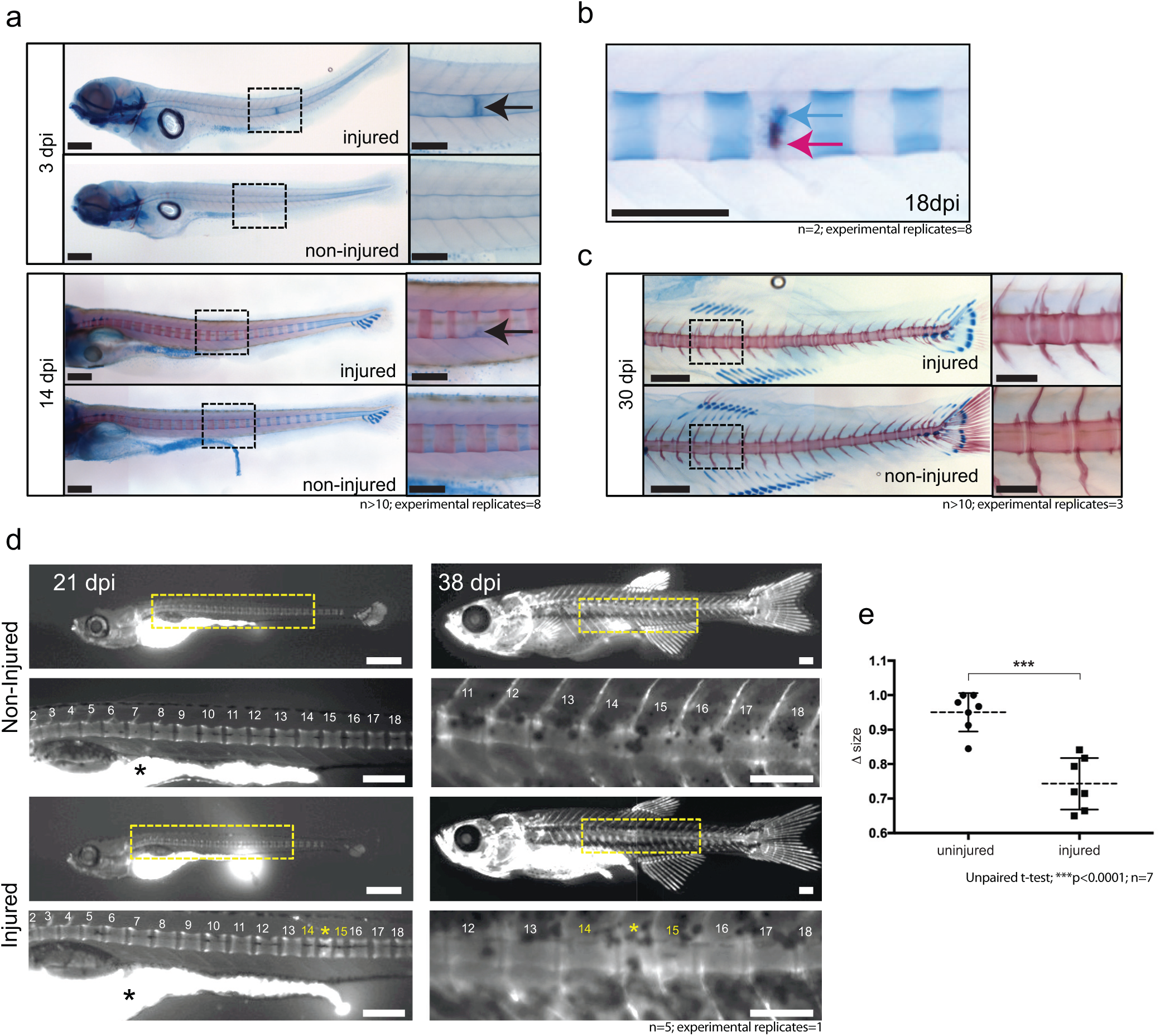
Ectopic vertebra formation occurs via a cartilage intermediate at the site of injury. (a) Alcian blue staining (cartilage staining) at the site of injury in 3 and 14 dpi larvae. Ectopic cartilage deposit is indicated by arrow. n >10; experimen-tal replicates = 8. Scale bar left panels: 400μm; scale bar right panels (zoomed images): 200μm. (b) Alcian blue and alizarin red (bone) staining at the site of injury 18 dpi indicates the presence of bone and cartilage at the repair site (blue arrow = cartilage; red arrow = bone). n = 2; experimental replicates = 8. Scale bar: 200μm. (c) Alcian blue and alizarin red staining of 30 dpi larvae reveals the formation of a smaller vertebra/vertebrae around the area of damage in injured larvae. n >10; experimental replicates = 3. Scale bar left panels: 400pm; scale bar right panels (zoomed images): 200ξm. (d) Live imaging of calcein stained zebrafish at 21 and 38 dpi in injured and uninjured fish. Extra vertebrae are indicated by (yellow asterisk). Black asterisk denotes intestinal fluorescence. n =5; experimental replicates = 1. Scale bar 21 hpf: 200μm; scale bar 21 hpf zoomed: 100μm; scale bar 38 hpf: 200μm; scale bar 38 hpf zoomed: 100μm. (e) The relative vertebra size difference (Δ size) between vertebrae at the site of injury (injured) and vertebrae in non-injured areas (uninjured). Vertebrae at the site of injury were significantly smaller than uninjured vertebrae (Unpaired t-test; ^∗∗∗^ p < 0.0001 two-tailed; mean +/- SEM uninjured larvae =0.9506 +/- 0.02102 n = 7; mean +/- SEM injured larvae =0.7432 +/- 0.0284 n = 7; measurements taken at 30 and 38 dpi).

By 18 dpi, the injured site began to express bone matrix, and was visibly flanked by cartilage expressing segments (**Figure 4b**). This was unusual because in development of the vertebrae, cartilage and bone stains mark distinct regions of the notochord. To evaluate the outcome of the injury in the bone process, wild-type larvae were injured and stained in calcein dye at 21 and 38 dpi (Du *et al*., 2001). Interestingly, the needle injury led to a delayed vertebral formation at the site of damage. These vertebrae that eventually formed were smaller and supernumerary, such that injured fish had one more vertebrae than their uninjured age-matched controls (**Figure 4 c-e**).

The notochord provides signals for the patterning of vertebral and spine formation via the patterned activation of various signals, and has been proposed to be an essential component of chordacentra formation (Flemming *et al*., 2004; Bensimon-Brito *et al*., 2012). The *Tg(entdp5:RFP)* transgenic line marks osteoblastic cells responsible for the patterned formation of chordacentra rings, and serves as a readout for mineralizing activity (Huitema *et al*., 2012). Entpd5 (ectonucleoside triphosphate diphosphohydrolase 5) is an E-type NTPase that is found in bone mineralizing environments and is essential for skeletal morphogenesis (Huitema et al., 2012). We crossed the *Tg(wt1b:GFP)* transgenic line to *Tg(entdp5:RFP)* and followed the wound response. *wt1b* and *entpd5* expressing cells populations were closely associated at the wound site indicating that mineralizing *entpd5* cells may directly contribute to *wt1b*+ associated chordacentra (**Figure 5 a,b**).

**Figure 5.**
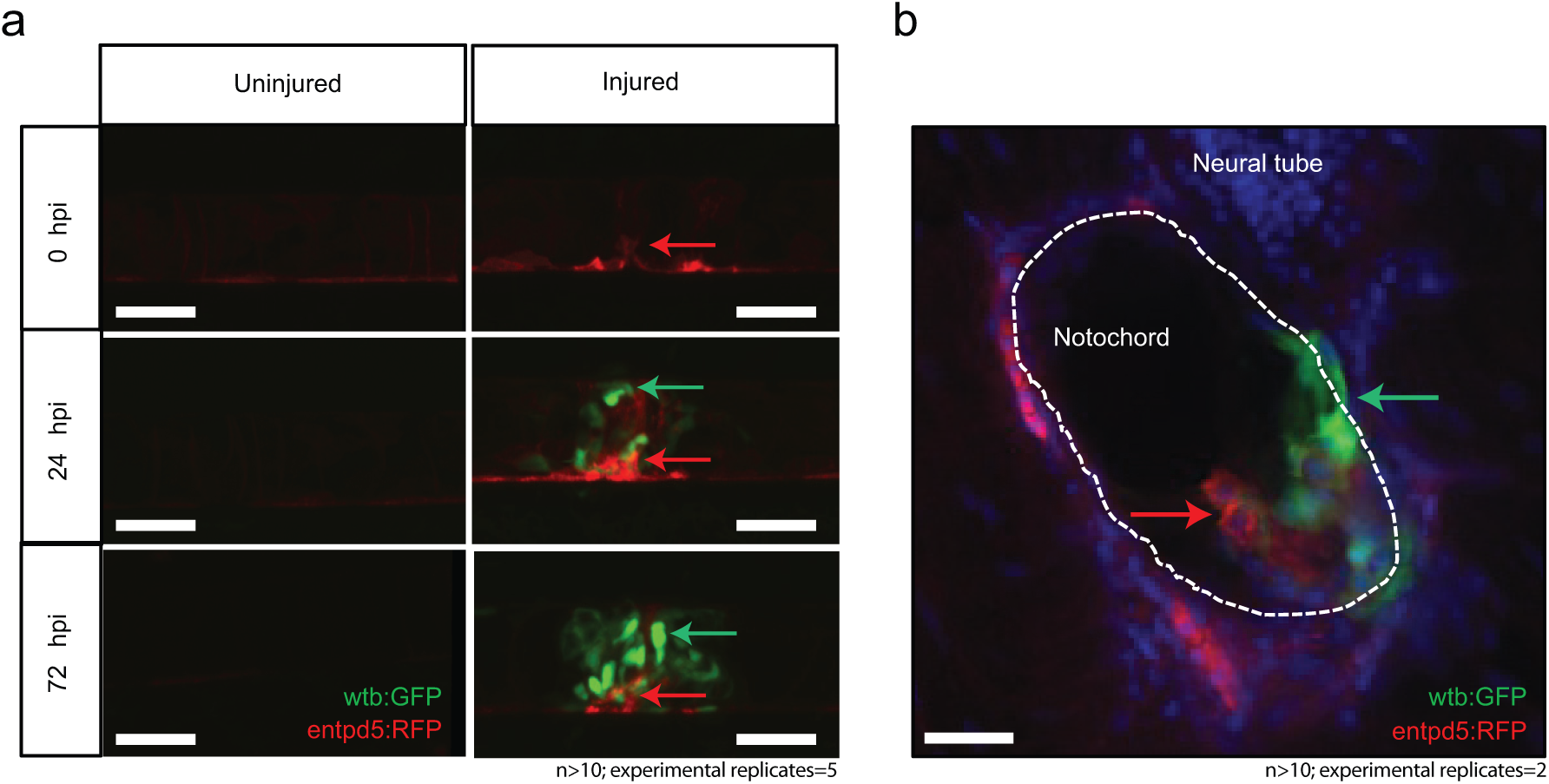
Distinct and closely associated wt1 and entpd5 subpopulations emerge at the damage site. (a) Live-imaging at the site of notochord injury in Tg(*wt1b:GFP; entpd5:RFP*) larvae. Expression of *wt1b:GFP* and *entpd5:RFP* at site of damage (green arrows and red arrows respectively) in injured and uninjured fish. n >10; experimental replicates = 5. Scale bar: 50μm. (b) Cryo-section of the injured area confirms distinct *wt1b:GFP* and *entpd5:RFP* subpopulations at site of damage. n >10; experimental replicates = 2. Scale bar: 20μm.

### Embryonic *wt1b*+ subpopulations perdure into the adult vertebrae

We noticed that the *Tg(wt1b:GFP)* transgene expression was always associated with the site of new vertebrae formation in the injured zebrafish that were raised to adulthood. To determine if *wt1b* expression was transient at the wound, or sustained throughout the repair process, we raised needle injured *Tg(wt1b:GFP; casper)* zebrafish larvae for up to 38 days (**Figure 6a**).

**Figure 6.**
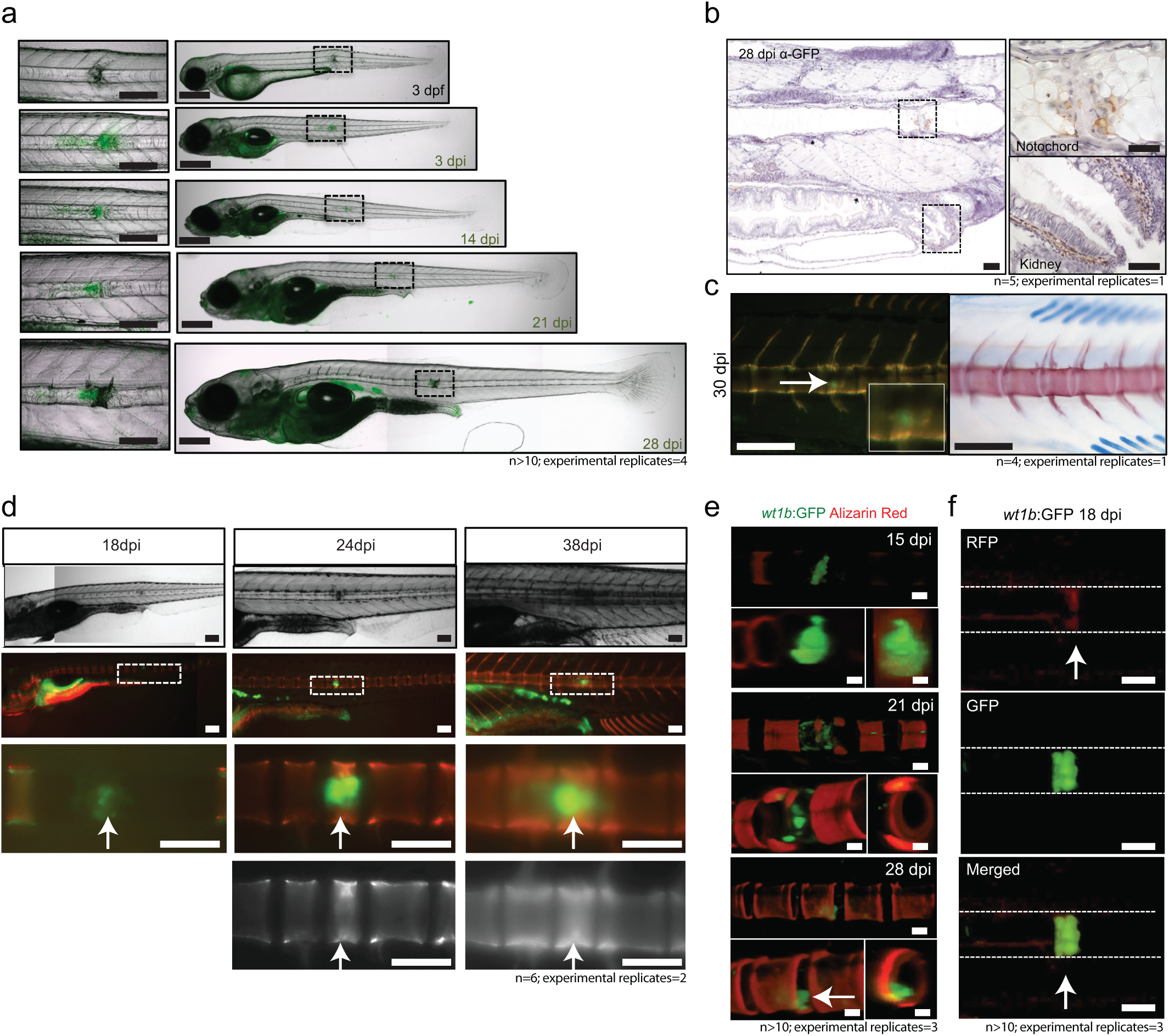
*wt1b* expressing cells are closely associated with vertebral development after injury. (a) Images of *wt1b:GFP* zebrafish following needle injury at 3dpf and raised to 28 dpi. n >10; experimental replicates = 4. Scale bar left panels: 100μm; scale bar right panels: 200μm. (b) α-GFP staining of 28 dpi larvae at the site of the healing notochord wound and in the kidney. n = 5; experimental replicates = 1. Scale bar left panels: 50μm. (c) Image of fish from Figure 5A, stained with alizarin red and imaged for *wt1b:GFP* expressing cells. GFP positive cells are found within the ectopic vertebra (white arrow and inset). n = 4; experimental replicates = 1. Scale bar left panels: 100μm. (d) Long term follow up of alizarin red stained Tg(*wt1b:GFP;casper*) larvae shows that chordacentra formation is delayed around the site of injury. GFP cells mark the site of the future ectopic vertebra. n = 6; experimental replicates = 2. Scale bar: 100μm; scale bar zoomed images: 50μm (e) Confocal imaging of 15, 21 and 28 dpi larvae reveals an overlapping expression between the wt1b:GFP expressing cells and the forming chordacentra (alizarin red stained) in the injured Tg(*wt1b:GFP;casper*) larvae. n >10; experimental replicates = 3. Scale bar: 100μm. (f) Confocal imaging highlights the overlapping presence of bone (alizarin red stained) and wt1b:GFP cells at the wound in 18 dpi larvae (arrow). n >10; experimental replicates = 3. Scale bar: 100μm.

GFP expression was sustained at the wound site, remaining in a small, cellular population at the site of damage, even as chordacentra developed and mineralized around the notochord over time (**Figure 6a, b, c**). Small GFP expressing cells were further confirmed by *α*-GFP staining at the site of damage (**Figure 6b**) Strikingly, the *Tg(wt1b:GFP)* transgene maintained expression at this site up to 38 dpi (**Figure 6d**) before eventually reducing expression.

To gain a better understanding of how *wt1b:GFP* expressing cells engage with the newly forming vertebrae, we carried out confocal imaging of the area of damage. The analysis revealed the presence of both fused and unfused vertebrae at the damaged site, and the sustained and strong expression of *wt1b:GFP* expressing cells associated with the developing ectopic vertebra at the repair site area (**Figure 6e,f**).

Taken together these results indicate that *wt1b:*GFP expressing cells both mark a subpopulation of cells that are rapidly activated at the site of the wound and also that these cells persist until adulthood, possibly orchestrating local vertebrae formation.

## Discussion

Our analysis has uncovered wound-specific cellular heterogeneity in the zebrafish notochord that perdures during adult vertebra formation at the injury site (**Figure 7**).

**Figure 7.**
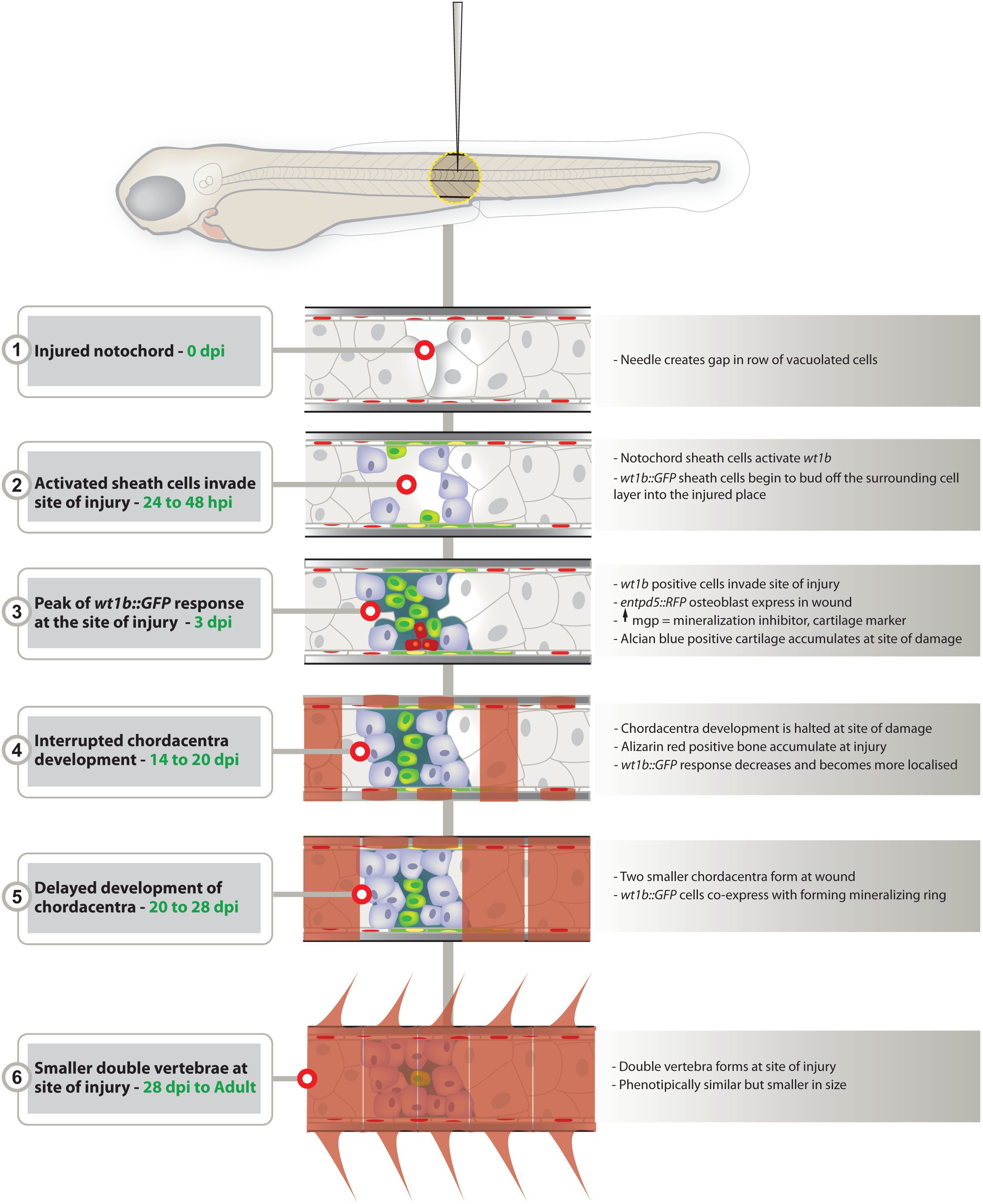
Schematic of the notochord wound response.

Despite *wt1b* having no reported role in notochord development, and despite not being expressed in the notochord, we identified a specific *de novo* expression of *wt1b* following notochord wounding. The activation of *wt1b* in sheath cells that migrate into the notochord is reminiscent of the situation where *wt1b* expression is reactivated in epicardial cells that undergo EMT to produce vascular progenitors and migrate into the heart (Martinez-Estrada et al., 2010). This raises the question whether notochord sheath cells may also be mesothelial in nature and if the invading *wt1b* expressing cells are produced via an EMT or, perhaps more accurately, a mesothelial to mesenchyme transition.

Wounding leads to localized *wt1b* expression in the notochord sheath cells that invade the site of the injury to form a stopper-like structure, likely to maintain notochord integrity. Very recently, Bagnet and colleagues reported the identification of notochord sheath cells involved in the replacement of vacuolated cells lost due to motion-dependent mechanical damage to the notochord (Garcia et al., 2017). In this context, sheath cells invade the vacuolated cell layer and differentiate into vacuolated cells to maintain turgor pressure. In light of this report, we have reanalysed our image analysis, but find no evidence to support that *wt1b* cells become vacuolated cells following acute wounding. In contrast, we find wt1b-expressing cells tightly associated with a stopper-like (scar-like) structure and continued *wt1b* expression at the wound site even during formation of an ectopic vertebra. We also detected *entpd5* expressing cell subpopulations at the wound that are distinct from *wt1b* expressing cells. These studies highlight a previously unknown complex and heterogeneous nature of the sheath, and suggest that the notochord sheath can sense and respond to different types of damage. Motion-dependent shear stress causes loss of vacuolated cells that are replaced by new vacuolated cells that arise from the sheath (Garcia et al., 2017), while acute damage (i.e. needle injury) that encompasses sheath and vacuolated cell damage, leads to sheath cells forming a seal that marks the site of new cartilage and vertebra (**Figure 7**).

By leveraging gene expression profiling of the wounded tissue, we discovered an alternative mechanism for vertebra formation via a cartilage intermediate at the injury site. This was unexpected because in zebrafish, ossification of chordacentra does not require the establishment of cartilage anlagen, but rather arises from the osteoblastic maturation of mescenchymal cells at the site of bone formation (Lefebvre & Bhattaram, 2010). Our observations indicate a wound-specific response to vertebra development. Vertebrae at the wound are supernumerary and smaller, with some showing defective neural and hemal arches (data not shown), and continue to be closely associated with *wt1b*+ cells until fully formed. We noted that the kinetics of vertebra formation at the damage site was delayed compared with other vertebrae. This delay could be explained by the very high expression of the cartilage gene *mgp* that inhibits calcification and BMP2 in mineralizing tissues (Schurgers *et al*., 2013; Zebboudj et al., 2002; Sweatt et al., 2003). This alternative mode for vertebra formation at the wound site may be a salvage structure to effectively maintain structural integrity of the developing axial skeleton.

## Materials and Methods

All experimental procedures were approved by the University of Edinburgh Ethics Committee and were in accordance with the UK Animals (Scientific Procedures) Act 1986.

### Zebrafish lines

Transgenic lines for this study include: *Tg(entpd5:RFP)* (Huitema et al., 2012), *Tg(ubi:switch)* (Mosimann *et al*., 2011), *Tg(SAGFF214:GFP)* (Yamamoto et al., 2010), *Tg(wt1a:GFP)* (Bollig et al., 2009), *Tg(wt1b:GFP)* (Perner et al., 2007; Bollig et al., 2009). Many of the studies were performed in a transparent background created by crossing homozygous *Tg(wt1b:GFP)* fish to homozygous pigment-free transparent *casper* fish (White et al., 2008). The *Tg(wt1b:GFP;col2a1a:RFP)* line was created by injecting the R2-col2a1a:mCherry construct (Dale and Topczewski, 2011) with a Tol2 transposase (Kawakami, 2007) into *Tg(wt1b:GFP;casper*) zebrafish embryos.

### Notochord needle injury and tail amputation assays

Larvae were anaesthetised in tricaine, placed sagittally on a petri dish and either inserted gently with an electrolysis-sharpened tungsten wire or tail amputated at different levels. Injured larvae were transferred to fresh water to recover and observe. Non-injured age-matched larvae were grown as non-injured controls.

### Whole-mount microscopy

Live and fixed whole-mount time-course and time-lapse experiments were performed using an AZ100 upright macroscope (Nikon) using a ×2 and ×5 lens with a Retiga Exi camera (Qimaging) or Coolsnap HQ2 camera (Photometrics). Images were analyzed and processed using the IPLab Spectrum and Micro-Manager softwares. Live and fixed whole-mount confocal imaging was performed using an A1R confocal system (Nikon) using a ×20 lens over a Z-plane range of 80-100 μm (approximate width of the notochord) using a 480nm laser (GFP) and/or a 520nm laser (RFP) lasers. Images were captured and analysed using Nis-Elements C software (Nikon). Multiphoton confocal time-lapse imaging was performed using an SP5 confocal microscope (Leica) equipped with a Ti:Sapphire multiphoton laser (Spectra Physics) and a 3 axis motorised stage. For confocal imaging and time-lapse experiments, anaesthetised injured and non-injured larvae were embedded sagittally in a drop of 1% low-melting point agarose prior to imaging, in a specially designed glass insert, which was covered in a mixture of E3 medium and anaesthetic. All time-lapse imaging was done at 30 or 60 mins intervals over 48 hours using an incubation chamber (Solent Scientific) under a constant temperature of 28°C and larvae were terminated in an overdose of tricaine at the end of each the experiment.

### Histology

Zebrafish larvae younger than 20 dpf were culled and fixed overnight in 4% PFA/PBS at 4°C. The fixed larvae were washed in PBS, dehydrated in rising methanol/PBS concentrations and cleared in xylene before being paraffin embedded for sectioning. Older zebrafish were culled and fixed in 4% PFA/PBS at 4°C for 3 days with an abdominal incision to ensure tissue penetrance of the fixative (Wojciechowska et al., 2016). Fish were decalcified using 0.5M EDTA (pH 7.5) for 5 days in a rocker at 4°C and dehydrated in 70% ethanol at 4°C. Fish were embedded in paraffin using a Miles Scientific Tissue TEK VIP automated processor. Embedded larvae and older zebrafish were sectioned using a Leica RM2235 rotary microtome to a width of 5 μm. Sections were haematoxylin and eosin (H&E) stained and mounted using DPX mounting media (Sigma-Aldrich). For cryosections, zebrafish larvae were embedded in OCT (Tissue Tek) and cut to 8 μm following protocols available at www.zfin.org.

### Immunohistochemistry

Slides were de-waxed in xylene and rehydrated through decreasing ethanol washes, before being incubated in a bleach solution to remove pigment. Antigen-unmasking was performed as previously described (Patton *et al*., 2005). The slides were DAB stained following manufacturer’s instructions (Dako). Slides were incubated overnight at 4°C with the following antibodies: *α*-GFP (1:1,500; Cell Signaling Technology) and *α*-WT1 (1:25,000). The *α*-WT1 was designed using the TARGET antibody production protocol from Cambridge Research Biochemicals using a conserved protein sequence from the C-terminal of the zebrafish Wt1a and Wt1b proteins. An Axioplan II fluorescence microscope (Zeiss) with a Plan Apochromat objective was used for brightfield imaging of tissue sections. Images were captured using a Qimaging Micropublisher 3.3mp cooled CCD camera and analysed using the IPLab Spectrum software.

### Immunofluorescence

Slides were processed as described above and blocked in 10% heat inactivated donkey serum for 2 hours. Slides were incubated overnight at 4°C with *α*-WT1 (1:33,000) antibody diluted in 1% heat inactivated donkey serum in TBSTw. Slides were incubated for 1 hour in a secondary anti-rabbit AlexaFluor 488 antibody (1:800) in 1% heat inactivated donkey serum and mounted in ProLong Gold mounting media containing DAPI overnight before being imaged in a fluorescent stereomicroscope.

### Tissue staining

Live bone staining was performed using 0.2% (w/v) calcein or using 50 μg/ml alizarin red as previously described (Du *et al*., 2001; Kimmel et al., 2010).

For cartilage and bone staining, alcian blue and alizarin red following the protocol outlined in (Walker and Kimmel, 2007) with modifications from protocols on www.zfin.org.

### RNA Extraction and microarray analysis

Fifty *Tg(wt1b:GFP)* zebrafish larvae were needle injured and grown to 72 hpi with age-matched non-injured controls. The area around the site of injury was dissected (Figure 4B) and transferred into 1 ml of chilled RNA-later. The samples were centrifuged into a pellet at 4°C and mascerated in 500 μl of Trizol^®^ (Sigma-Aldrich) using a 25G ^5/8^ 1 ml syringe. RNA was extracted following Trizol^®^ manufacturer’s instructions and eluted into 15 μl of distilled H_2_O. Extracted RNA was sent to Myltenyi Biotec (Germany) who conducted the microarray analysis. Injured and non-injured samples were sent in triplicates and the RNA was amplified and Cy3-labelled using a Low Input Quick Amp Labelling Kit (Agilent Technologies) following manufacturer’s instructions. The labelled cRNA was hybridised against a 4×44K Whole Zebrafish (V3) Genome Oligo Microarray (Agilent Technologies). The microarray images were read out and processed using the Feature Extraction Software (FES – Agilent Technologies) and differential gene expression was determined using the Rosetta Resolver^®^ gene expression data analysis system (Rosetta Biosoftware).

### Fluorescence-Activated Cell Sorting

The trunk region of fifty *Tg(wt1b:GFP; R2-col2a1a:RFP)* injured larvae and non-injured 72 hpi larvae were dissected and collected separately in cold PBS+2% fetal calf serum (FCS). Tissue disassociation was adapted from a previously described protocol (Manoli and Driever, 2012) and centrifuged cells were collected in FACSmax cell disassociation solution (Genlantis). The samples were passed twice through a 40 μm cell strainer, collected in an agar-coated petri dish on ice and transferred into an eppendorf tube to be sorted by a FACSAria2 SORP instrument (BD) equipped with a 405nm, a 488nm and a 561nm laser. Green fluorescence was detected using GFP filters 525/50BP and 488nm laser, red fluorescence was detected using 585/15BP filter and 561nm laser. Data was analyzed using FACSDiva software (BD) Version 6.1.3.

### Vertebrae size measurements and statistical analysis

The vertebrae size difference in injured zebrafish larvae (age range 30 dpi to 38 dpi) were compared between vertebrae at the site of injury (injured) and vertebrae outside of the site of injury (uninjured). Injured vertebrae and uninjured vertebrae were measured and the average length was recorded for each group. The average lengths were then compared and the relative size difference was calculated. The relative size difference between each group (injured:uninjured vs. uninjured:uninjured) was compared using an unpaired t-test.

## Acknowledgements

We thank Craig Nicol for assistance with figure design, Andrea Coates for critical reading of the manuscript, and the Zebrafish Facility staff at the MRC Human Genetics Unit for zebrafish husbandry.

## Video 1 Legend

Time-lapse imaging of two-photon microscopy of *Tg (wt1b:GFP; col2a1a:RFP)* zebrafish larvae following needle injury over 48 hours. *wt1b:GFP* expression is upregulated in *col2a1a:RFP* expressing notochord sheath cells upon needle injury, leading to the formation of a stopper like structure across the wound

